# Adaption of a Conventional ELISA to a 96-well ELISA-Array for Measuring the Antibody Responses to Influenza virus proteins, viruses and vaccines

**DOI:** 10.1101/2019.12.20.885285

**Authors:** Eric Waltari, Esteban Carabajal, Mrinmoy Sanyal, Natalia Friedland, Krista M. McCutcheon

## Abstract

We describe an adaptation of conventional ELISA methods to an ELISA-Array format using non-contact Piezo printing of up to 30 spots of purified recombinant viral fusion proteins, vaccine and virus on 96 well high-protein binding plates. Antigens were printed in 1 nanoliter volumes of protein stabilizing buffer using as little as 0.25 nanograms of protein, 2000-fold less than conventional ELISA. The performance of the ELISA-Array was demonstrated by serially diluting n=8 human post-flu vaccination plasma samples starting at a 1/1000 dilution and measuring binding to the array of Influenza antigens. Plasma polyclonal antibody levels were detected using a cocktail of biotinylated anti-human kappa and lambda light chain antibodies, followed by a Streptavidin-horseradish peroxidase conjugate and the dose-dependent signal was developed with a precipitable TMB substrate. Intra- and inter-assay precision of absorbance units among the eight donor samples showed mean CVs of 4.8% and 10.8%, respectively. The plasma could be differentiated by donor and antigen with titer sensitivities ranging from 1 × 10^3^ to 4 × 10^6^, IC_50_ values from 1 × 10^4^ to 9 × 10^6^, and monoclonal antibody sensitivities in the ng/mL range. Equivalent sensitivities of ELISA versus ELISA-Array, compared using plasma and an H1N1 HA trimer, were achieved on the ELISA-Array printed at 0.25ng per 200um spot and 1000ng per ELISA 96-well. Vacuum-sealed array plates were shown to be stable when stored for at least 2 days at ambient temperature and up to 1 month at 4-8°C. By the use of any set of printed antigens and analyte matrices the methods of this multiplexed ELISA-Array format can be broadly applied in translational research.

## INTRODUCTION

The activity of humoral antibodies provide the best correlation to long-term immune memory and protection (Antia et al. 2018). During the first two weeks of exposure to a pathogen, the majority of antibodies found in the serum derive from plasmablasts, either rapidly re-activated from memory B cell pools or expanded from newly stimulated, somatically hypermutated and differentiated B cells upon contact with antigen in lymph tissue (De Silva and Klein 2015). During recovery, some plasmablasts will home to the bone marrow where they terminally differentiate into long-lived plasma cells stably secreting antibodies that circulate in serum for many years (Abbas, Lichtman, and Pillai 2014; Yoshida et al. 2010). Cellular and molecular events leading to antigen-specific B cell expansion, differentiation, homing and fate are complex and not predictable in outcome. In lieu, serum can be used to measure the binding kinetics, magnitude, specificity and cross-reactivity of the secreted antibodies in response to infection or vaccination. Serological testing can help evaluate an individual’s susceptibility, exposure or protection from past, existing and future pathogens. It is also possible to make positive or negative correlations of binding characteristics to serum neutralization activity or antibody enhanced disease (Katzelnick et al. 2017). Analytical methods characterizing antibodies ideally have the ability to measure the robustness, specificity and genetic breadth of activity to pathogens. Humoral responses are typically quantified by titer in naïve, acute, convalescent and recovery sera in the context of natural infection or pre- and post-vaccination and correlated to *in vitro* activity assays and clinical signs of immune protection (e.g. Antia et al., 2018; Lowell et al., 2017; Madore et al., 2010).

The enzyme-linked-immunosorbent-assay (ELISA) first described by Engvall and Perlmann (1972), is commonly used to measure specific antibody-antigen binding. Variants and derivatives of the ELISA have become assay workhorses of immunology laboratories and a host of compatible reagents, consumables, plate washers, multichannel pipettes, robotic liquid handlers, and assay formats have been developed and are available from multiple vendors. A conventional antigen ELISA single plex format passively coats antigens on a 96-well high capacity protein binding surface (e.g. Nunc Maxisorp™, ThermoFisher Scientific, Waltham, MA) and indirectly titers primary antibody binding by secondary binding of polyclonal antibodies conjugated to horseradish peroxidase (HRP), which turns over the colorimetric 3,3’,5,5’-tetramethylbenzidine (TMB) substrate for assay readout. Secondary antibodies are typically directed against a constant region of the heavy or light chain of the primary antibody, such as polyclonal anti-Fc directed to IgG, IgM, IgA or IgE, anti-kappa or anti-lambda light chains. A common variation to boost sensitivity includes using a biotinylated secondary antibody with a Streptavidin-horseradish peroxidase (HRP) conjugate. Although fluorescent reporters have the advantage of allowing for multiplexed detection using different dyes, the use of HRP has been shown to be more sensitive because the enzymatic turnover of colorimetric or chemiluminescent substrates amplifies the signal (Gogalic et al. 2018).

The principles of ELISA have been adapted using advances in the protein array field to increase the throughput, efficiency and scope of data in immunoassays (reviewed in Kingsmore 2006). Printing proteins can be carried out by passive adsorption without requiring modification or chemical coupling to nanoparticles or other surfaces. This advantage and advancements in nozzle technology allow for flexibility and precision in spotting picolitre volumes of purified, crude, or complex proteinaceous substrates (Barbulovic-Nad et al. 2006). Furthermore, a superior level of sensitivity can be achieved in miniaturized ligand-binding assays, as shown by Ekins’ ambient analyte assay theory (Ekins 1989). Obtaining higher sensitivity in a system that uses smaller amounts of capture molecules and smaller amounts of sample can be explained by two main features. First, the binding reaction occurs at a high target concentration; and second, the capture-molecule–target complex is found only in the small area of the spot, resulting in a high local signal (Templin et al. 2002). The most published format for protein array printing in the infectious disease research setting is onto glass slides functionalized with nitrocellulose, perhaps because both of the technical ease and that high density arrays are made possible by printing onto this high protein binding flat surface (Davies et al. 2005; Desbien et al. 2013; Koopmans et al. 2012; Nakajima et al. 2018; Price et al. 2013; te Beest et al. 2014). An alternative format amenable for use in research labs is an ELISA-based microarray printed directly onto the bottom of a 96-well plate (Mendoza et al. 1999). This method has been validated against single plex assays (Liew et al. 2007) and has been adopted for biomarker discovery in research labs (W. Huang et al. 2018; Y. Huang and Zhu 2017) and commercial assays (e.g. PBL Assay Science, Piscataway, NJ; Quantarix, Billerica, MA; BioVendor LLC, Asheville, NC; RayBiotech Inc., Peachtree Corners, GA). However, to date the 96-well format has been infrequently applied to infectious disease antigens (Kang et al. 2012; Wang et al. 2015), warranting more published examples and methods of applied research in this area.

For our multiplexed infectious disease research, we decided to capitalize on the resources, familiarity and knowledge readily available for the conventional 96-well plate ELISA and adapt the workflow directly to an in-house 96-well plate ELISA-Array format. The only changes in the assay format were at the first and final steps. Using Maxisorp™ 96-well plates in the first step, in lieu of coating a single antigen per well, we printed 1 nanoliter volumes of 8 viral antigens, in triplicate, per well. In the final step, a precipitating form of TMB substrate and an array plate reader were needed for the ELISA-Array instead of the soluble TMB form and general lab plate reader. The remainder of the workflow and reagents were identical in both formats. The development and testing of the ELISA-Array was carried out using healthy human donor plasma sampled post-vaccination (from the 2018 FluLaval vaccine), and assayed on printed vaccine, recombinant hemagglutinin trimers, and purified FluB virus. Here we provide data characterizing the ELISA-Array methods, advantages, precision, robustness, sensitivity, stability and utility in infectious disease research.

## RESULTS

### Initial optimization of printing conditions

Although many operating conditions for printing followed the standard recommendations of the manufacturer of the sciFLEXARRAYER S12 instrument, several specific parameters were optimized for this ELISA-Array application. We tested variations in printing protein concentration, drop volume and formulation buffer using goat anti-human Fc polyclonal antibody (Jackson ImmunoResearch, West Grove, PA) as a probe with commercially available human reference serum (Bethyl Laboratories, Montgomery, TX) as an analyte. The probe was varied by diluting a PBS stock in a 1:1 volume of each of three sciSPOT protein formulation buffers D1, D11 and D12 (Scienion AG, Berlin, Germany). Probe was dispensed in 1, 2 or 4 drops from a 384-well source plate at 25, 100 and 400 ug/mL final. PBS in formulation buffers without the anti-human Fc protein was used for a background control. The probes of the printed arrays bound to the Fc region of IgG within the human plasma, then are detected with HRP conjugate antibodies specific to the kappa light chain of the IgG antibody in a traditional sandwich ELISA format.

The signal intensity increased with increasing protein printed, and 400 ug/mL provided the highest signal. The spot size increased with drop number, but the sensitivity was similar between 2 and 4 drops. The protein stabilizing D12 buffer offered the highest sensitivity among formulation buffers to approximately 4 ng/ml concentrations of IgG detected from human sera. These data are shown in Supplementary Figure S1. The final printing parameters used in this report for Influenza antigens are described in the methods section. Twelve 96-well plates were printed in one batch with the Influenza antigens listed in Table 1 and using the array pattern illustrated in Figure 1.

**Table 1.** Antigens included in the protein microarrays.

| Influenza Strain | Subtype |
| --- | --- |
| A/Michigan/45/2015* | H1N1 |
| A/Japan/305+/1957 | H2N2 |
| A/Singapore/INFIMH-16-0019/2016* | H3N2 |
| A/Viet Nam/1194/2004 | H5N1 |
| A/Shanghai/02/2013 | H7N9 |
| B/Phuket/3073/2013-like virus* (B/Yamagata/16/88 lineage) | Influenza B |
| B/Colorado/06/2017-like virus* (B/Victoria/2/87 lineage) | Influenza B |
| Flulaval Vaccine 2018-2019 | *H1N1, *H3N2,<br>*Influenza B -1, *Influenza B -2 |

\* components of the WHO recommended seasonal flu vaccine for 2018-19

**Figure 1.**
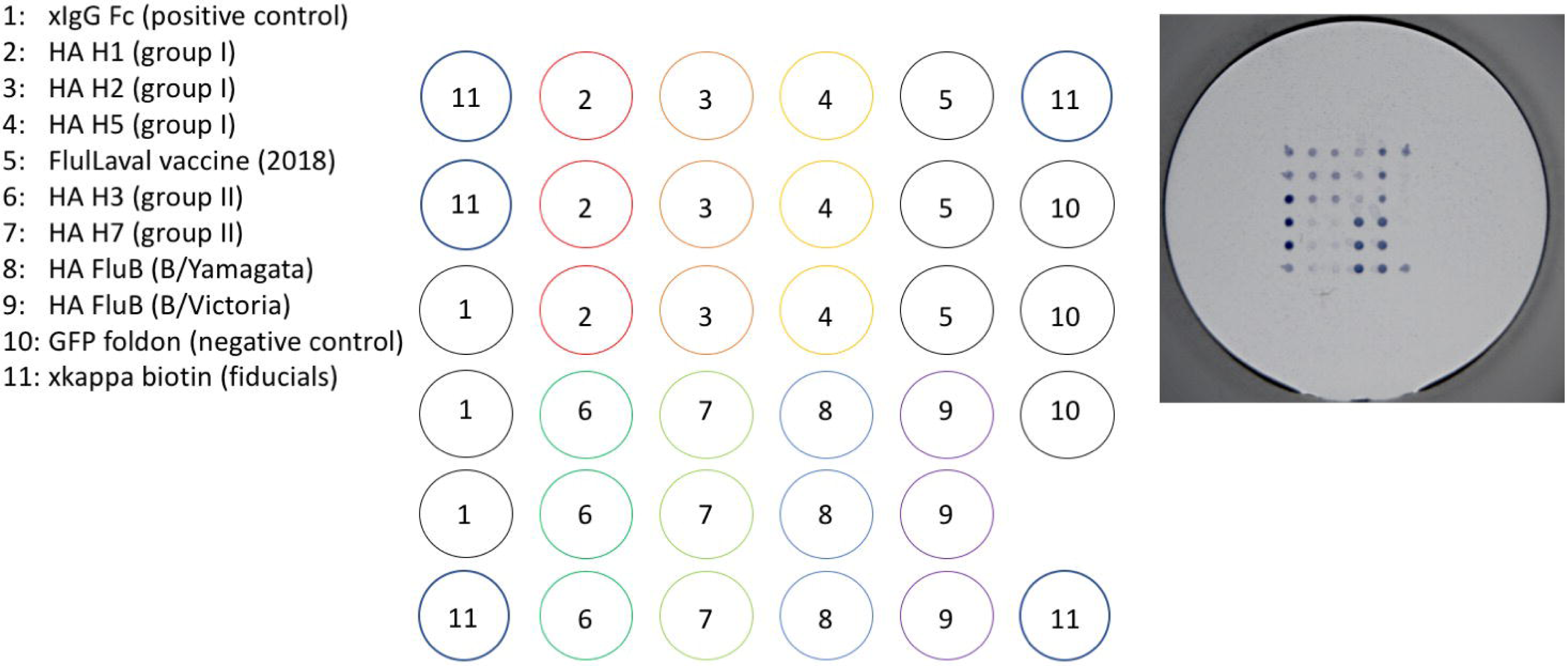
The ELISA-Array print pattern. The 6×6 array was printed in each well of a 96-well plate. The outer edges contain fiducial spots of biotinylated anti-kappa antibody for orientation (#11). All other spots are printed in triplicate, using anti-IgG Fc antibody as a positive control (#1) and GFP-foldon as a negative control (#10). In the top half of the pattern are Influenza A HA group I proteins (#2-4) and the 2018 vaccine (#5), and in the bottom half are Influenza A HA group II proteins (#6-7) and Influenza B HA proteins (#8-9). At right is an image of one array well developed after binding reference plasma on H1N1 A/Michigan/45/2015 HA trimer.

### Assay miniaturization gain of sensitivity in ELISA-Array

According to the ambient assay theory (Ekins 1989), miniaturizing the ELISA to an array print of 0.25ng of protein in a 200um spot (with a surface area of 15.6mm^2^, or 0.02ng/mm^2^) should yield higher sensitivity than coating in the same proportion over an entire 96-well (with a surface area of 320mm^2^). We tested this by comparing the signal sensitivity obtained using human immune reference plasma binding purified H1N1 HA trimer, either printed in 0.25ng spots in triplicate or coated in a 96-well at 1000, 100, 10 or 1 ng in duplicate. Following the same assay methods with the exception of the final TMB substrate (soluble for the ELISA and precipitating for the ELISA-Array), equivalent IC_50_ values were obtained only when the 96-well was coated with 2000-fold more total protein, or >150-fold more/mm^2^ (Figure 2).

**Figure 2.**
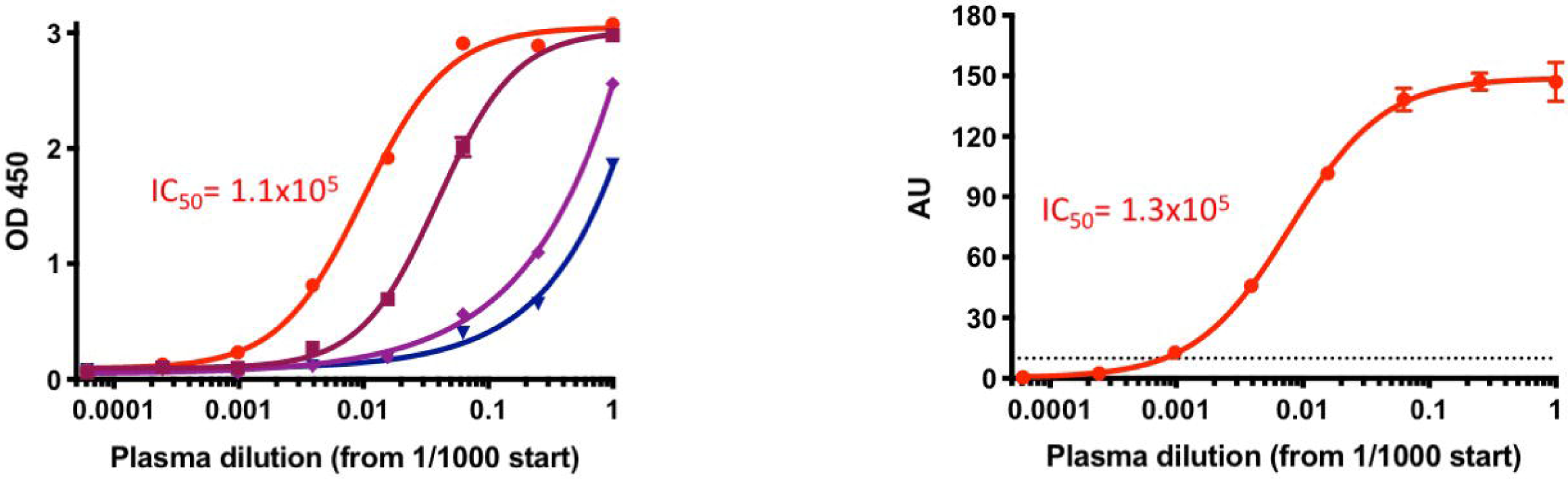
A comparison of the amount of protein needed in ELISA versus ELISA-Array. Two sets of data are shown from a titration of a reference plasma on the H1N1 A/Michigan/45/2015 HA trimer. On the left, a conventional ELISA is shown using 4 different antigen coat amounts. On the right, ELISA-array data is shown for the same antigen printed at 0.25ng where an equivalent IC_50_ is obtained when compared to the highest (1ug) antigen coat on the conventional ELISA. The dashed line in ELISA-array plot indicates the lower limit of quantification (10 AU).

### Data analysis

After calculating median intensity in absorbance units (AU) of each triplicate set of antigen spots, we fit standard 4-parameter logistic (4P) curves of intensity against plasma or mAb concentration with PRISM (GraphPad, San Diego, CA). In each plate we tested three negative controls to calculate the lower limit of detection, and on average the LOD value was less than 5 AU. Because the variance in readings at values less than 10 AU was high (data not shown), we set a lower limit of detection (LLOQ) at 10 AU. From the 4P curves fit to each sample we calculated both the titer at which the curve passed the LLOQ and the IC_50_ values. Across our assays we observed that the upper intensity range was never greater than 180 AU and thus set limits of 0-200 AU in 4P curve fitting. Because at high concentrations the hook effect can lead to reduced intensity readings (Tighe et al. 2015), we disregard any decreased values at high analyte concentrations.

### ELISA-Array Sensitivity and Specificity

In each ELISA-Array assay, polyclonal human immune reference plasma and monoclonal antibodies of known binding activity were used to control for assay performance and determine sensitivity. The dose-dependent binding of each of these controls over three independent assays (Figure 3), IC_50_ values and LLOQ are reported in Table 2. mAb A is known to be a neutralizing antibody recognizing a conformationally dependent epitope on the stalk region of HA trimers (Kallewaard et al. 2016) and was detected against the array of Influenza antigens from 10-120ng/mL, well below the quantitative ug/mL range of relevant protective antibody levels *in vivo* (Crum□Cianflone et al. 2012; Semenova et al. 2004). Reference plasma showed Influenza antigen binding IC_50_ values of 1.4 × 10^5^ to 9 × 10^6^ and titers of 6.4 × 10^4^ to 1 × 10^6^ (Figure 3 and Table 2). These values reflect a polyclonal mixture of antibodies of any isotype since detection was not limited to IgG (a cocktail of anti-kappa and anti-lambda light chain secondary antibodies was used). The correlation of binding titers to protection varies by disease and for Influenza has not been shown to be predictive (Madore et al. 2010). However, since binding is a pre-requisite for neutralization activity, plasma titer can demonstrate the variation, breadth and magnitude of viral antigen specificity between individuals.

**Table 2.** Sensitivity of reference mAbs and reference plasma pAb on ELISA-array antigens.

| Antigen | IC<sub>50</sub> (ng/mL) | LLOQ (ng/mL) | Plasma titer IC<sub>50</sub> | Plasma titer LLOQ |
| --- | --- | --- | --- | --- |
| mAb A |  |  | Reference plasma |  |
| Vaccine | 1900 | 470 | 1.4e5 | 2.6e5 |
| H1N1 | 110 | 10 | 8.5e6 | 1.0e6 |
| H2N2 | 380 | 120 | 1.3e5 | 2.6e5 |
| H3N2 | 170 | 30 | 4.0e5 | 2.6e5 |
| H5N1 | 40 | 10 | 3.0e5 | 2.6e5 |
| H7N9 | >30,000 | >30,000 | 2.9e5 | 6.4e4 |
| mAb B |  |  |  |  |
| Vaccine | 7.3 | 2 |  |  |
| Influenza B1 | 238 | 125 | 9.0e6 | 1.0e6 |
| Influenza B2 | 25 | 2 | 4.6e6 | 1.0e6 |

**Figure 3.**
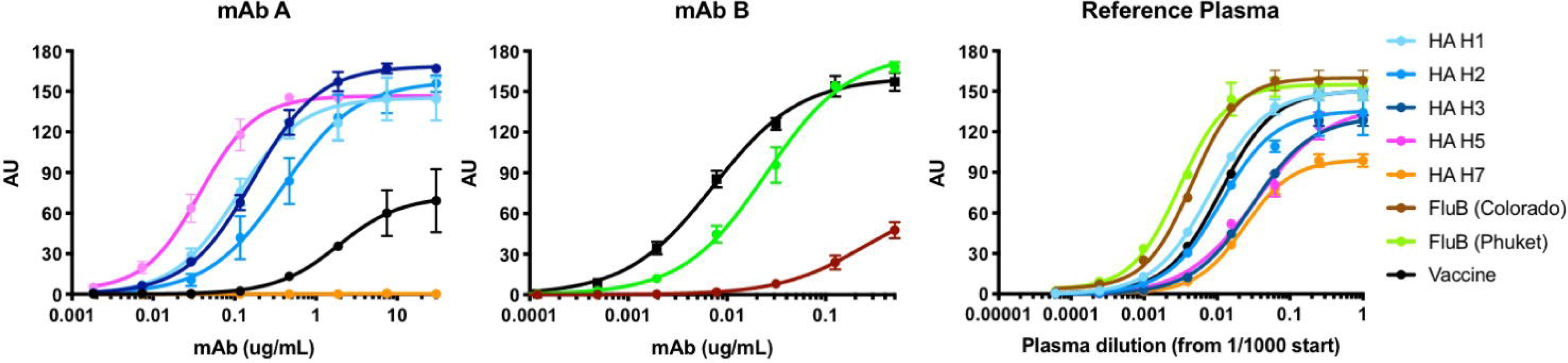
The sensitivity of the ELISA-Array. Three sets of data are shown with the titration of two monoclonal anti-Influenza antibodies and a reference plasma on the Influenza array antigens. Panels show titrations of an anti-Influenza A mAb (left), an anti-Influenza B mAb (center) and a reference plasma (right) against the array of Influenza A antigens (left & right), Influenza B antigens (center & right), and the FluLaval 2018-2019 vaccine (all). Data is from 3 inter-assay plates, with mean and SD shown.

The specificity of the assay was tested by measuring cross-reactivity at 200nM concentrations of two irrelevant mAbs to any of the printed proteins: one directed to the envelope protein of Dengue virus and the other to the RSV fusion protein. No signal was observed in the assay with these mAbs. Specificity was also tested by printing a GFP-foldon-Avitag-6His trimer in each well as a negative control protein, at the same concentration as the Influenza A HA trimers. This control showed no binding to donor plasma or to control antibodies mAb A and mAb B.

The lack of binding of control mAb A to the HA trimer of A/Shanghai/02/2013 H7N9 was not expected based on publications of this mAb binding to other H7 strains of HA, albeit at lower affinities than other HA subtypes (Kallewaard et al. 2016). Reference plasma and other donor plasma did bind the H7 trimer (Figures 3 and 4). A repeat test print of the H7 HA trimer at 0.5, 0.375 and 0.25ng/spot did not change the binding results, nor did testing on a regular ELISA format (data not shown). Further optimization of this antigen, and comparisons to other strains are needed in order to draw conclusions on cross-reactive antibody binding to H7.

**Figure 4.**
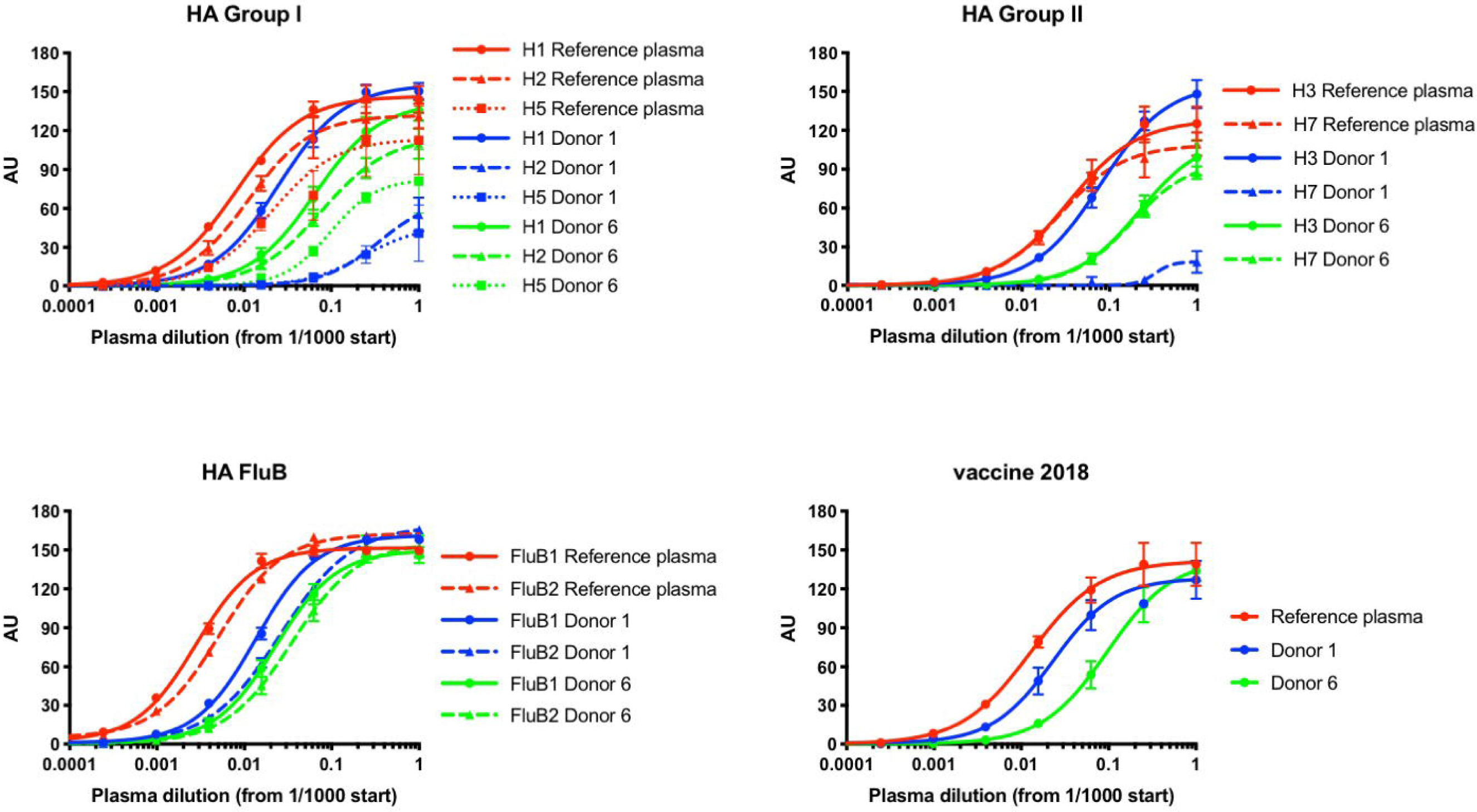
Dose-response curves of 3 donors on the Influenza antigen array. The dose-dependent binding curves of a reference plasma (red), donor 1 plasma (blue) and donor 6 plasma (green) to the Influenza antigen array are shown. The top left panel shows plasma titrations against Influenza A group I HA trimers, the top right panel shows titrations against Influenza A group II HA trimers, the bottom left panel shows titrations against Influenza B viruses, and the bottom right panel shows titrations against the FluLaval 2018-2019 vaccine. Data is from 3 inter-assay plates, with mean and SD shown.

### ELISA-Array assay performance

The ELISA-Array assay was qualified using a selected in-house human reference plasma and eight individual human plasma samples from day 28 post-vaccination with the 2018 FluLaval quadrivalent vaccine (GlaxoSmithKline, Research Triangle Park, NC). All array plates used for performance testing were from one print batch, stored at 4°C. To increase accuracy, we avoided making large dilutions by preparing stock solutions of 10x reference plasma, 10x control mAbs, 100x secondary biotinylated antibody mixture, and 100x streptavidin conjugate in assay diluent. These were aliquoted and stored at -80°C. Each plasma donor was also aliquoted undiluted and stored at -80°C. Although not done in these assays, it would be optimal in the future to briefly spin down donor plasma before assaying to clear the sample of lipid and other aggregates. Aliquots were freshly thawed for each assay, and the same lot of assay diluent and TMB substrate were also used throughout all assays. The final concentrations of assay materials are described in the methods section. Intra-assay precision was determined by running n=3 plate assays on the first day after printing the arrays. Inter-assay precision was determined by running an additional two plates one and two weeks later. Precision was calculated by the variance between plates of titer and IC_50_ values for reference plasma and each of the eight donors for all array antigens. Intra- and Inter-assay precision data is shown for the reference plasma and two donors in Figure 4, and Tables 3 and 4, respectively, and for all eight plasma samples in Supplementary Tables S1 and S2. The precision of absorbance units among the reference plasma and two donors showed mean CV of 4.8% intra-assay and 10.8% inter-assay, and 6.0% intra-assay and 12.5% inter-assay among the eight donor samples. There were a few examples of high variance inter-assay, in samples of low dilutions. This may be due to weak binding or interference from the serum matrix. The plasma titers could be differentiated by donor and antigen with sensitivities ranging from 1 × 10^3^ to 4 × 10^6^ and IC_50_ values from 1 × 10^4^ to 9 × 10^6^ (Figure 4, Table 4, and Table S2). For example, we measured a robust titer in the reference plasma donor to all array antigens (Figure 4). In contrast, robust titers in donor 1 plasma were measured only to the vaccine itself, to Influenza B viruses and to the individual antigens of H1N1 and H3N2 HA trimers matching the strains used in the vaccine (Table 1 and Figure 4). There was only weak binding to HA trimers not in the vaccine (i.e. H2, H5, H7), indicating insufficient cross-reactive antibodies were elicited in donor 1. A plot of IC_50_ values in Figure 5 for three donors shows the overall tight standard deviations between three assays performed over three weeks, and visually quantifies differences between antibody binding for each of the array antigens and donors. We cannot differentiate pre-existing antibody immunity from vaccine responses in these samples, but the quantitative nature of the data would allow for this to be done using titer and IC_50_ comparisons with pre-vaccine plasma, not included in this study. Three rounds of freeze thaws of the reference plasma from -80°C showed no change in IC_50_ values or titers (data not shown). One plate assay was also run by a second operator to evaluate the robustness of the method, which was determined to be equivalent to intra-assay precision (Plate 4 in Figure 6 and Table S3).

**Table 3.**
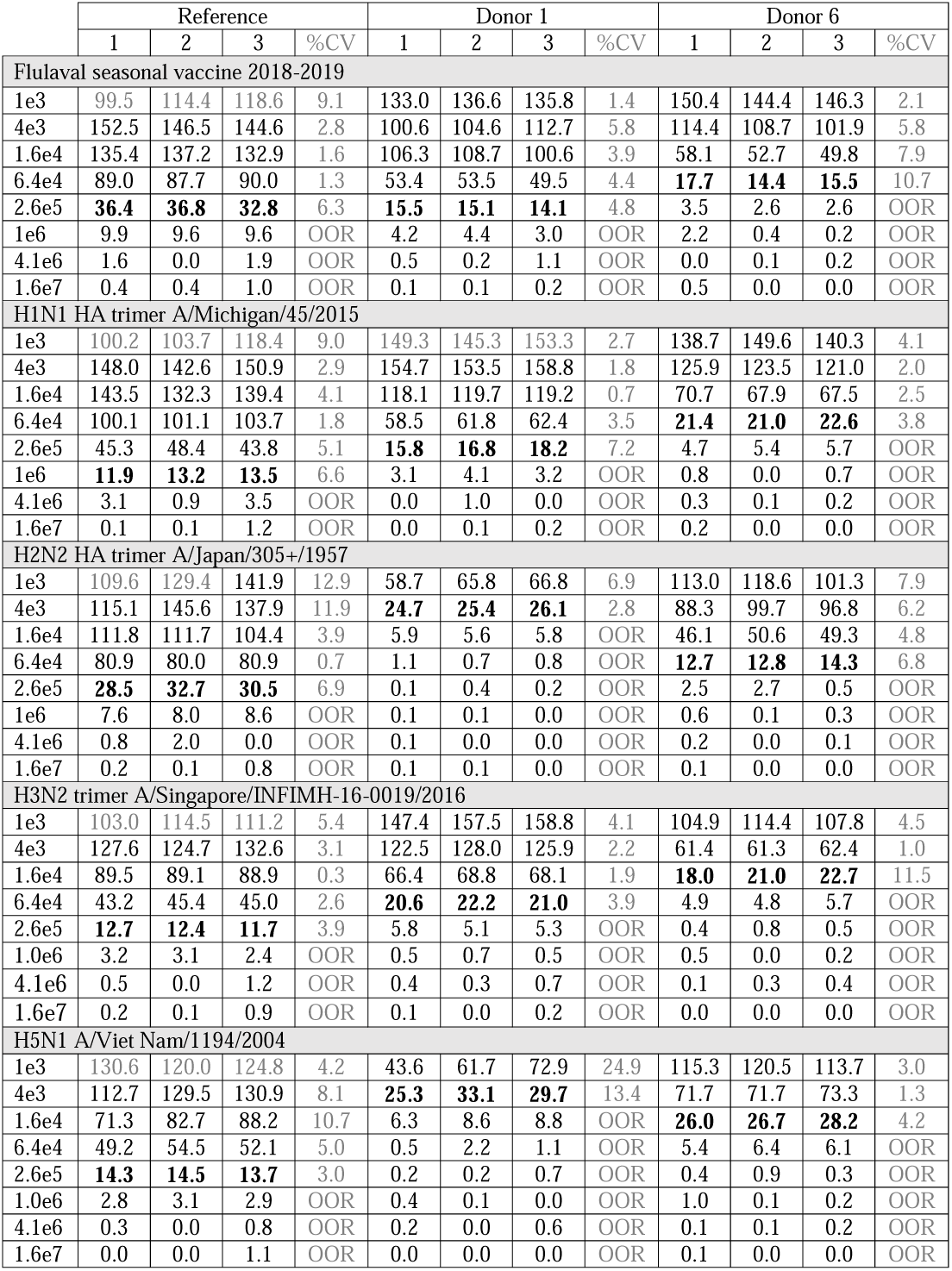
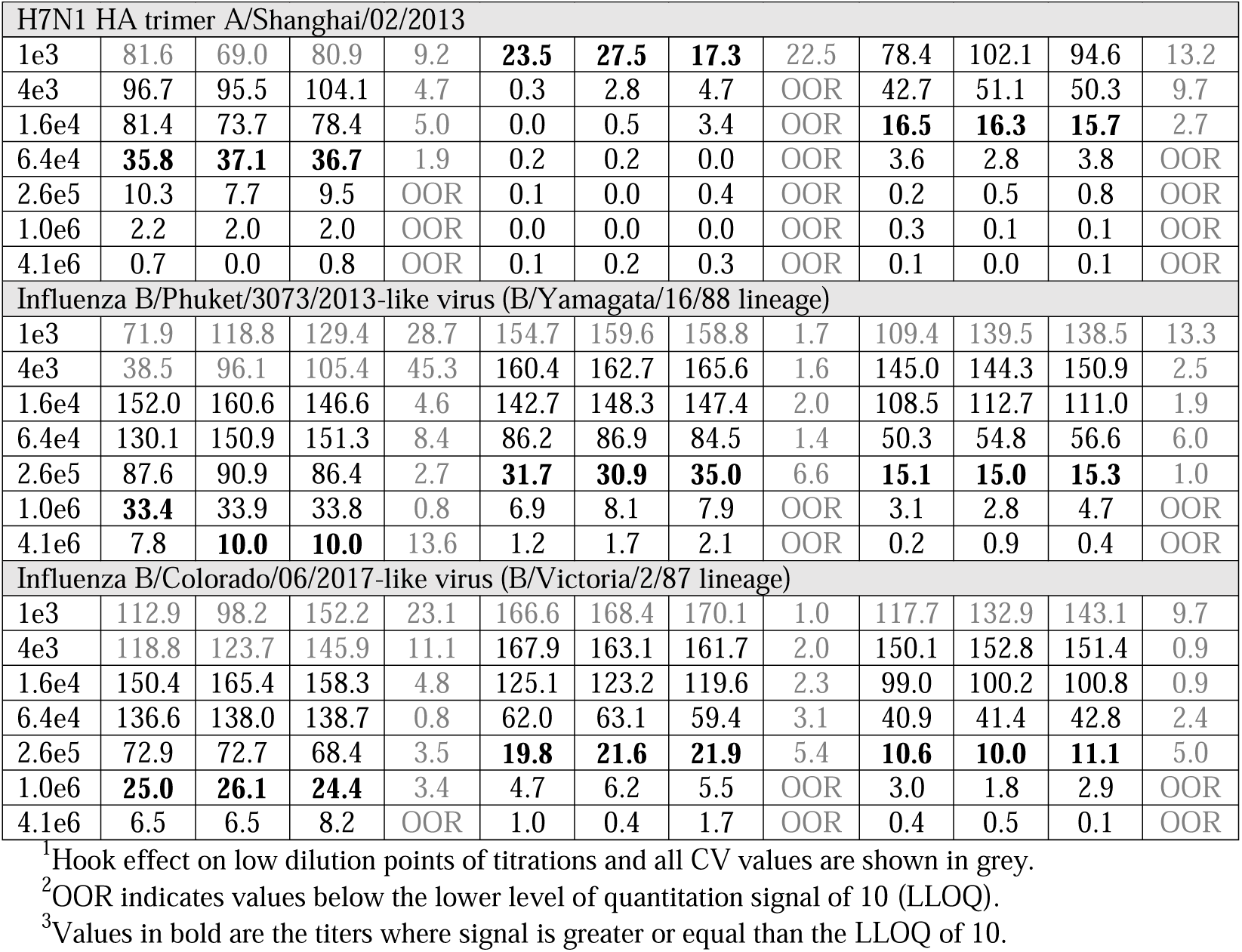
Intra-assay precision from three donor plasma samples on ELISA-array antigens.^1, 2, 3^

**Table 4.**
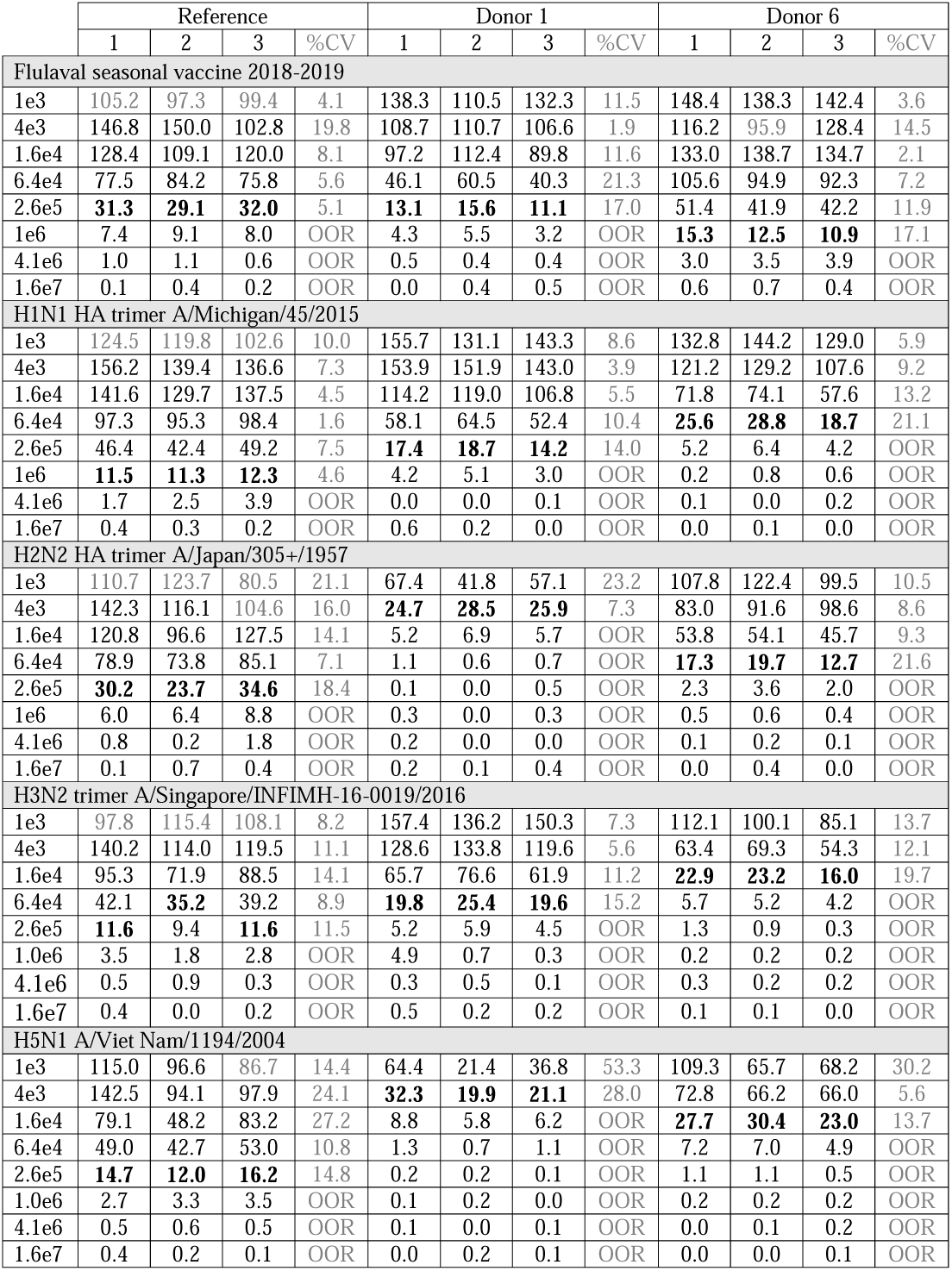
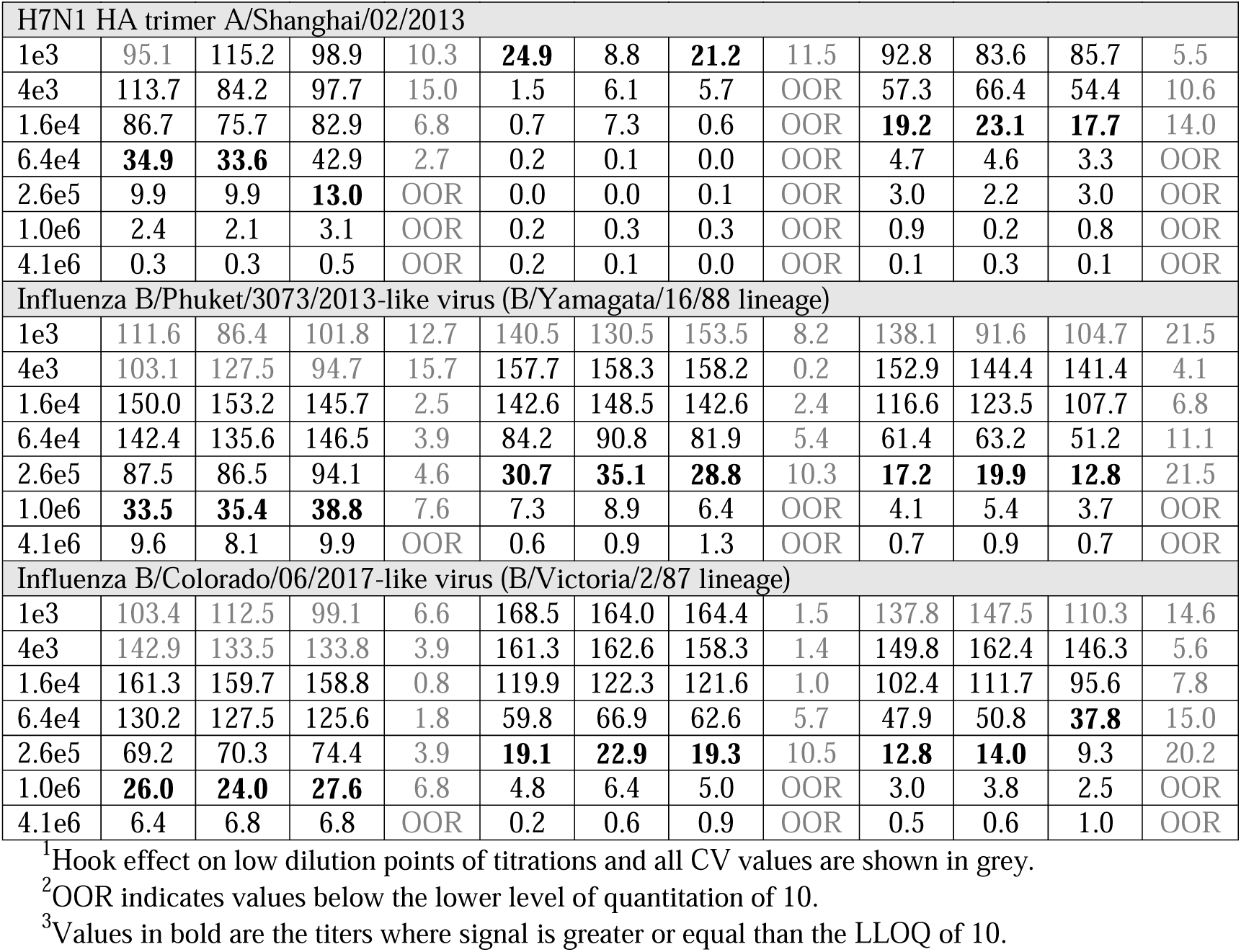
Inter-assay precision from three donor plasma samples on ELISA-array antigens.^1, 2, 3^.

**Table 5.** Intra-assay and Inter-assay precision of plasma IC_50_ values on ELISA-Array antigens.

|  | Reference |  |  |  | Donor 1 |  |  |  | Donor 6 |  |  |  |
| --- | --- | --- | --- | --- | --- | --- | --- | --- | --- | --- | --- | --- |
|  | 1 | 2 | 3 | %CV | 1 | 2 | 3 | %CV | 1 | 2 | 3 | %CV |
| Flulaval seasonal vaccine 2018-2019 |  |  |  |  |  |  |  |  |  |  |  |  |
| Intra | 1.2e5 | 1.1e5 | 1.1e5 | 6.0 | 2.2e5 | 2.2e5 | 2.6e5 | 10.1 | 1.1e4 | 1.2e4 | 1.6e4 | 20.3 |
| Inter | 1.5e5 | 1.4e5 | 1.4e5 | 4.8 | 3.0e5 | 1.4e5 | 3.4e5 | 40.5 | 1.3e4 | 7.1e5 | 1.2e4 | 28.0 |
| H1N1 HA trimer A/Michigan/45/2015 |  |  |  |  |  |  |  |  |  |  |  |  |
| Intra | 8.6e6 | 7.3e6 | 8.6e6 | 8.8 | 2.24e5 | 2.0e5 | 2.2e5 | 6.3 | 6.4e5 | 8.1e5 | 7.1e5 | 11.8 |
| Inter | 1.0e5 | 8.4e6 | 7.1e6 | 17.6 | 2.6e5 | 2.4e5 | 2.6e5 | 4.1 | 5.6e5 | 6.2e5 | 8.1e5 | 20.0 |
| H2N2 HA trimer A/Japan/305+/1957 |  |  |  |  |  |  |  |  |  |  |  |  |
| Intra | 8.8e6 | 1.0e5 | 1.2e5 | 16.7 | 5.1e4 | 5.6e4 | 5.5e4 | 5.3 | 1.1e4 | 8.3e5 | 6.4e5 | 24.8 |
| Inter | 1.4e5 | 1.3e5 | 1.2e5 | 9.2 | 6.0e4 | 1.7e4 | 3.8e4 | 55.3 | 6.6e5 | 1.1e4 | 6.6e5 | 29.9 |
| H3N2 trimer A/Singapore/INFIMH-16-0019/2016 |  |  |  |  |  |  |  |  |  |  |  |  |
| Intra | 4.0e5 | 3.6e5 | 4.6e5 | 13.5 | 8.1e5 | 8.7e5 | 9.4e5 | 7.3 | 2.6e4 | 3.7e4 | 3.0e4 | 18.7 |
| Inter | 4.6e5 | 4.1e5 | 3.4e5 | 14.9 | 9.1e5 | 5.4e5 | 9.7e5 | 29.1 | 3.3e4 | 1.7e4 | 2.0e4 | 36.3 |
| H5N1 A/Viet Nam/1194/2004 |  |  |  |  |  |  |  |  |  |  |  |  |
| Intra | 5.2e5 | 5.2e5 | 4.2e5 | 12.0 | 2.4e4 | 3.0e4 | 7.6e4 | 66.0 | 2.3e4 | 2.7e4 | 2.0e4 | 14.4 |
| Inter | 5.6e5 | 1.9e5 | 1.4e5 | 75.8 | 3.6e4 | 9.3e5 | 2.6e4 | 56.2 | 1.9e4 | 9.8e5 | 7.9e5 | 48.4 |
| H7N1 HA trimer A/Shanghai/02/2013 |  |  |  |  |  |  |  |  |  |  |  |  |
| Intra | 2.4e5 | 2.6e5 | 3.0e5 | 13.0 | >1e3 | >1e3 | >1e3 | OOR | 3.9e4 | 4.2e4 | 3.5e4 | 8.2 |
| Inter | 3.2e5 | 3.7e5 | 1.9e5 | 30.6 | >1e3 | >1e3 | >1e3 | OOR | 2.3e4 | 1.2e4 | 2.1e4 | 31.6 |
| Influenza B/Phuket/3073/2013-like virus (B/Yamagata/16/88 lineage) |  |  |  |  |  |  |  |  |  |  |  |  |
| Intra | 3.3e6 | 3.3e6 | 3.2e6 | 2.3 | 1.4e5 | 1.4e5 | 1.6e5 | 7.1 | 3.2e5 | 2.2e5 | 3.2e5 | 20.1 |
| Inter | 3.1e6 | 3.3e6 | 2.1e5 | 112.0 | 1.4e5 | 1.3e5 | 2.5e5 | 39.1 | 2.7e5 | 2.0e5 | 2.0e4 | 124.0 |
| Influenza B/Colorado/06/2017-like virus (B/Victoria/2/87 lineage) |  |  |  |  |  |  |  |  |  |  |  |  |
| Intra | 4.2e6 | 5.2e6 | 4.5e6 | 11.4 | 2.6e5 | 2.7e5 | 3.0e5 | 7.7 | 2.3e5 | 2.8e5 | 3.2e5 | 15.7 |
| Inter | 4.5e6 | 5.7e6 | 3.8e6 | 21.3 | 2.9e5 | 2.3e5 | 2.5e5 | 10.0 | 2.7e5 | 3.8e5 | 5.1e5 | 30.6 |

**Figure 5.**
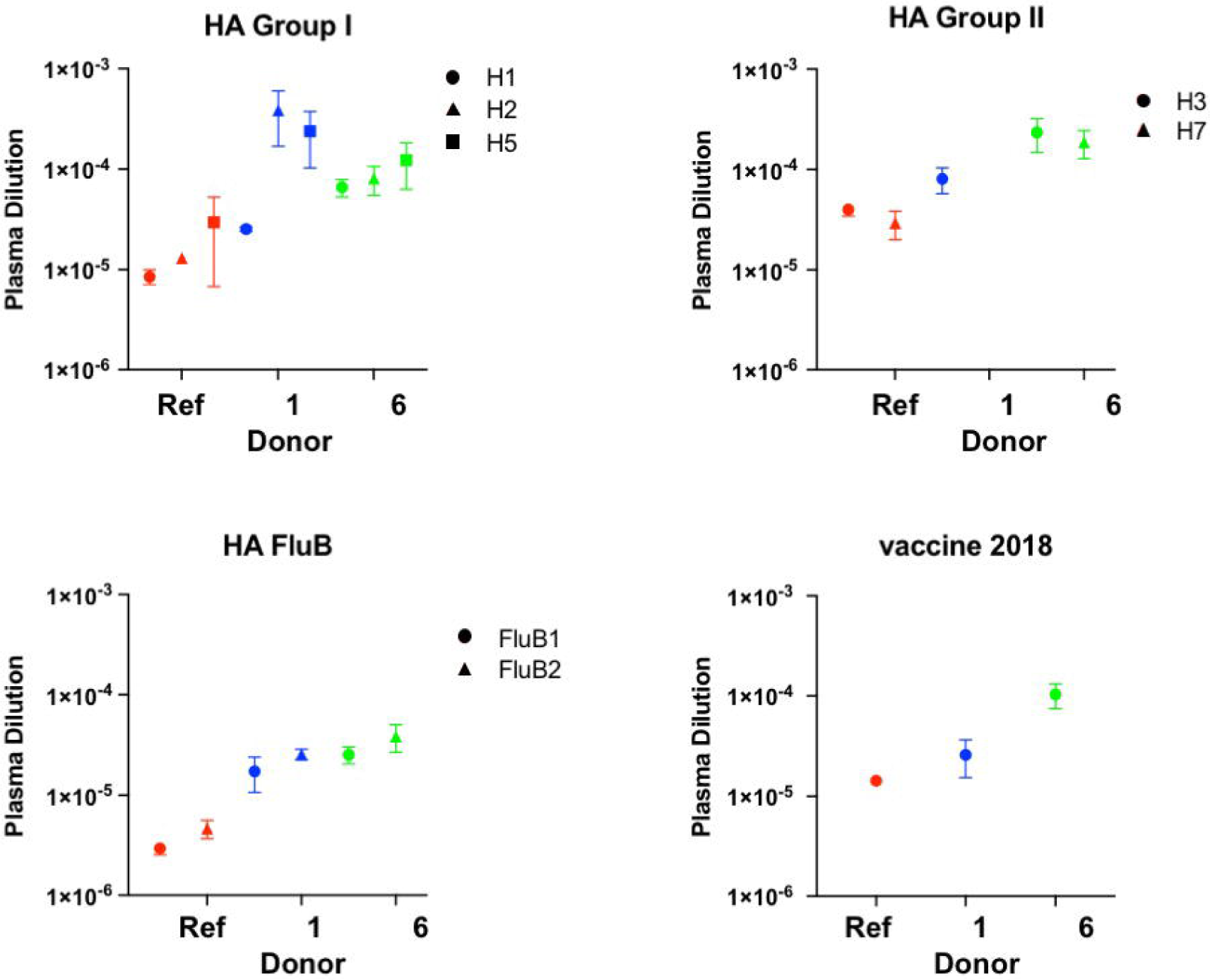
IC_50_ comparisons of 3 donors on the Influenza antigen array. IC_50_ values determined from 4-parameter fits of array antigen binding curves using a reference plasma (red), donor 1 plasma (blue) and donor 6 plasma (green) are plotted. The top left panel shows IC_50_ values against Influenza A group I HA trimers, the top right panel shows IC_50_ values against Influenza A group II HA trimers, the bottom left panel shows IC_50_ values against Influenza B viruses, and the bottom right panel shows IC_50_ values against the FluLaval 2018-2019 vaccine. Data is from 3 inter-assay plates, with mean and SD shown. The IC_50_ value for donor 1 plasma against H7 is not shown because the titer was greater than the minimum 1/1000 dilution.

**Figure 6.**
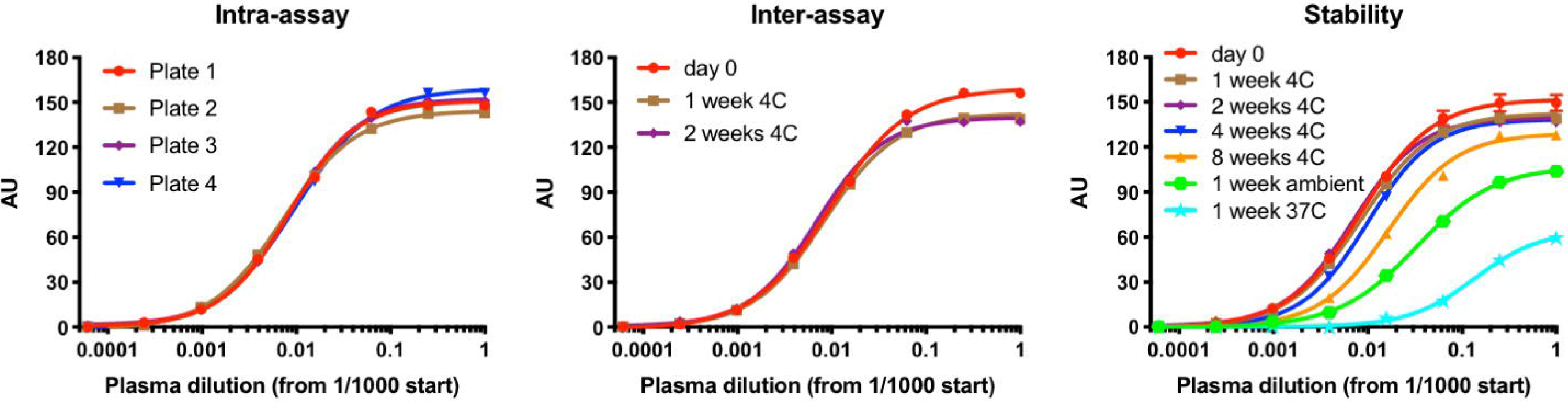
Intra-assay, Inter-assay and Stability Performance of the ELISA-Array. Three dose-dependent binding curves are shown using a reference plasma and the H1N1 A/Michigan/45/2015 HA trimer. The left panel shows both intra-assay error (plates #1-3) as well as inter-operator robustness (plates #1-3 vs. #4). The center panel shows inter-assay error with plates tested after increasing time periods stored at 4 °C. The right panel shows stability after 1 week at ambient temperature, at 37 °C and up to 8 weeks at 4 °C.

### Stability testing of printed plates

Printed array plates were covered with a foil plate seal and vacuum-sealed immediately after the overnight curing step and stored at 4°C. They were found to be stable stored in this manner for up to one month (Figure 6). At 8 weeks post-print, plates stored at 4°C showed about a 2-fold drop in IC_50_ and the titer shifted to one higher dilution in the ¼ titration series (i.e. a change from 1×10^6^ to 2.6 ×10^5^). Significant losses in activity were also measured for vacuum-sealed plates stored for 1 week at either ambient temperature or 37 °C (Figure 6). All of the stability assay data including variability for each antigen and plasma sample are provided in Supplementary Tables S4 and S5. Using a different print lot, we tested the plates for 2-days at ambient temperature and found them to be stable (data not shown).

## DISCUSSION

An ability to print microarrays in a format for a 96-well ELISA-Array was first published by Mendoza et al. (1999), and its utility for infectious disease testing has been demonstrated with antibody arrays to encephalitis viruses (Kang et al. 2012) and viral antigen arrays to Flaviviridae (Wang et al. 2015), using a non-contact piezo Bio-Dot Printing System (Biodot, Irvine, CA). As with these two prior studies, we printed using non-contact piezo nozzles, but in smaller volumes using a Scienion S12 instrument. We compared binding data in arrays using a variety of Influenza antigen types including recombinant viral protein HA trimers, vaccine and viruses. The parallel identification of viral antigen binding was carried out in a quantitative manner by performing full dose-response curves of human plasma with an analysis of the precision of titer and IC_50_ values. These data allowed for a comparison of the abundance and context of antibodies from natural exposure or vaccination within a single individual and between individuals. In future arrays it would be of interest to include the Influenza neuramidase protein, a second viral surface protein that can be targeted by neutralizing antibodies (Memoli et al. 2016). It is important to note that the quality and relevance of protein array data is only as good as the proteins printed. The use of reference monoclonal antibodies or plasma with known activity is helpful to characterize the integrity of the protein reagents and printing conditions. Such reference reagents can also serve to bridge data between different array print lots and stability as well as between data from different operators or labs. Stability testing of the ELISA-Array plates supports the ability to ship plates on cold packs to a collaborating research lab, with a tolerance of up to 48 hours at ambient temperature. The collaborating lab would only incur the lesser cost and training required for the array reader.

There is great potential to use what is learned from protein arrays in research labs towards the design and testing of vaccines and for the development of simplified rapid point-of-care testing (POCT) of infectious diseases. To date, POCT efforts have taken the form of printed arrays on lateral flow test strips, a promising technology needing further development to be useful in endemic, low resource settings (Kim, Chung, and Kang 2019; Urusov, Zherdev, and Dzantiev 2019). Protein array data can also be a valuable tool for the identification of individuals most likely to have acquired broadly cross-reactive and/or potent neutralizing antibodies to infectious disease. Passive immunization with monoclonal antibodies can be highly effective in controlling viral pathogenesis (Salazar et al. 2017), and an ability to screen the serum of human donors suspected to be clinically protected from disease in a multiplexed and quantitative format can help identify the best donor for antibody discovery. In this application it is valuable to have functional neutralization assay data on the same sera, to correlate to binding.

Overall, we have provided new methods and qualification data to support applications ELISA-Array assay format for infectious disease research. The key advantages we observed with this technology included the passive coating in a protein stabilizing buffer, the low protein reagent consumption with nanoliter printing, and the ability to perform quantitative analyses using nearly the same workflow, reagents and lab equipment as used in the conventional ELISA.

## METHODS

### Chemicals

Phosphate buffered saline (Gibco DPBS, calcium and magnesium free, pH 7.2) and EDTA (0.5M Ambion) were obtained from Thermofisher (Waltham, MA). PBS with 0.05% Polysorbate-20 was purchased as a 20x stock from Teknova Inc. (Hollister, CA). Fraction V bovine serum albumin (BSA), gamma-Globulins from bovine blood (BGG), ProClin 300 and CHAPS were purchased from Sigma-Aldrich (St. Louis, MO). Fetal Bovine Serum (FBS), US source, triple 0.1um filtered, was sourced from Omega Scientific (Tarzana, CA).

### General instrument and experimental parameters

The Scienion sciFLEXARRAYER S12 instrument (Scienion AG, Berlin, Germany) has been optimized for non-contact, piezo-acoustic dispensing of ultra-low volumes from an inert coated glass capillary in a climate-controlled (temperature, dewpoint and humidity) environment, with precise XYZ axis control and on-board camera and software for QC of each spot in the array.

We printed our arrays with the PDC70 type 3 nozzle (Scienion) due to its reduced dispense volume and the specific hydrophobic coating optimized to improve the dispense stability of protein solutions. The system liquid is Milli-Q Water filtered through the Milli-Q Ultrapure Water System (Millipore Sigma, Burlington, MA) that is subsequently degassed for at least 30 minutes in a sonicating water bath. Proteins were spotted on low dust, black, clear bottom, high protein binding Fluotrac™ 600, Greiner Bio-One 96-well plates (Fisher Scientific, Waltham, MA), positioned on a ceramic platform under vacuum. The printing was carried out at 60% humidity and ambient temperature, with drop volumes of 330-350 pL. The 384-well source plate (Scienion) was kept at dewpoint. Drop stability and array quality were assessed for quality for each run. Prior to dispensing into the plates, autodrop detection was used to assess drop stability by quantifying the velocity, deviation and drop volume for each protein spotted. In addition, all plates were imaged with the on-board head camera after the completion of spotting to ensure correct alignment and spot diameters. Printed arrays were incubated overnight at 75% relative humidity and ambient room temperature to allow adsorption of the proteins to the binding surface of the plate. Plates were then vacuum packaged and stored at 4°C until ready for use.

The sciREADER CL2 (Scienion) is used for the colorimetric reading of the final assay of arrays. After images are taken of each well, the software analysis program aligns the spot pattern to the imaged spots and calculates a median intensity in absorbance units (AU).

### Initial ELISA-Array assay used for optimization of parameters

Initial 96-well printed arrays were printed according to the general instrument parameters described above. All assay steps were performed at ambient temperature. Blocking solution (sciBLOCK Protein D1M solution, Scienion) was added at 200 μL/well with a multichannel pipet and allowed to incubate without agitation for 1 hour. The block solution was manually removed and the plate washed 1x by adding 300 μL/well of sciWASH Protein D1 agitating at 350rpm for 5 minutes on a Bioshake iQ thermomixer. Dilutions of human reference serum (with IgG quantified at 4.4 mg/ml; Bethyl Laboratories) were made in blocking buffer (PBS, 0.05% Tween-20, and 0.5% BSA), and 100 μL/well was added to the plate incubated for 1 hour with gentle agitation (250 rpm). The arrays were manually washed 3 times with 2-5 min agitations in between washes. A secondary polyclonal goat antibody (Southern Biotech, Birmingham, AL) directed to the human kappa light chain and conjugated with horseradish peroxidase (HRP) was used at 1/5000 in blocking buffer for 1 hour. After a second round of 3 manual washes the signal was developed with sciCOLOR T2, a precipitating TMB reagent (Scienion), for 15 minutes.

### Influenza hemagglutinin proteins

Recombinant hemagglutinin (HA) ectodomain constructs were made using gene blocks (IDT, Newark, NJ) cloned into a pADD2 plasmid using EcoRI and XhoI restriction sites. The cloning was performed with In-Fusion HD Cloning kit (Takara Bio, Mountain View, CA). Each construct consisted of the native HA signal sequence, HA ectodomain, a trimeric foldon domain of T4 fibronectin, an Avi tag sequence (GLNDIFEAQKIEWHE), and a hexa-histidine affinity tag (Whittle J. et al., 2014). A negative control construct was made with the tags fused to the GFP protein. Plasmids were purified with NucleoBond® Xtra Maxi kit (Macherey Nagel, Düren, Germany) and transfected into Expi293 cells grown in a 1:2 mixture of Expi293 and Freestyle (Gibco, Thermofisher Scientific) media. For transfection, 50 μg of plasmid was pre-incubated with 1.3 ml of FectoPRO transfection reagent (Polyplus, New York, USA) and added to 1L of media. At day 4, the media was clarified by centrifugation (7500xg, 15 min), filtered, and diluted 2-fold with PBS. The media was then batch incubated with HisPur Ni-NTA resin (Thermofisher Scientific) for 2 hours at 4°C and loaded on a gravity flow column. The resin was washed with 20 column volumes of PBS with 5 mM imidazole and eluted in 4 ml of PBS with 250 mM imidazole. The eluted protein was concentrated and loaded on a Superdex 200 16/60 column (GE Healthcare, Chicago, IL) pre-equilibrated with PBS. The elution fractions corresponding to the trimeric HA proteins were pooled, concentrated, and stored in 10% glycerol in PBS at -20°C.

### Human plasma and reference antibodies

Post-vaccination (28 days) blood samples were collected from nine healthy volunteers who received the 2018/2019 seasonal influenza vaccine (FluLaval quadrivalent vaccine, GlaxoSmithKline, Research Triangle Park, NC) in the fall of 2018 following a protocol approved by Stanford University (IRB protocol 48130). Plasma was separated from heparinized blood samples by centrifugation at 500g for 15 minutes and stored at -20°C. One donor was identified with the highest plasma titer to all Influenza antigens and was used as a reference in all assays. Two specific positive control reference antibodies with characterized low nanomolar affinities to HA trimers were cloned and made recombinantly as human IgG1 in Expi293 cells with methods described previously (Durham et al. 2019): MEDI8852 for InfA group 1 and 2 HA (Kallewaard et al), and TF19 for InfB HA (unpublished in-house reagent). Two recombinant mAbs were used as cross-reactivity controls, J9 directed to the Dengue viral envelope (Durham et al. 2019) and 3D3 to the RSV fusion protein (Collarini et al. 2009).

### Printing of the Influenza protein array

Protein stocks of recombinant proteins at 0.5 mg/mL in PBS or vaccine stocks were diluted 1:1 in D12 buffer, mixed by pipetting, transferred to a 384 well polypropylene plate (sciSOURCEPLATE, Scienion), and centrifuged for 2 min at 1800xg ambient temperature to eliminate debris or air bubbles. The pattern printed was a 6×6 spot array on each well, and each protein or vaccine along with positive and negative controls was printed in triplicate, 3 spots per well, with 3 drops printed per spot. A single lot of twelve 96-well plates were printed in one day, and after overnight curing were either subject to the ELISA-Array assay the next day (plates 1-4), subject to temperature variations for one week (plates 5-7), or subject to varying incubation times at 4°C (plates 8-10).

### Influenza ELISA-Array assay

Each 96-well printed array was printed according to the general instrument parameters described above. All assay steps were performed at ambient temperature, incubations except the blocking step were done with low agitation on a Titer Plate Shaker (Lab-line Instruments, Melrose, IL) and washes were done using a BioTek ELx405 plate washer (Bio-Tek Instruments, Winooski, VT). High agitation is avoided as it leads to comets around the spots, which interferes with accurate spot definition during reading of absorbance intensity. Array plates were washed 1x 300 μL/well before immediately adding 200 μL/well of assay diluent (PBS, 0.5% BSA, 2% filtered FBS, 0.2% BGG, 0.25% CHAPS, 5mM EDTA, 0.05% Polysorbate-20 and 0.05% ProClin 300, pH 7.2) down the sides of the wells with a multichannel pipette. Plates were allowed to block for 1 hour. Human plasma was diluted 1/1000 in assay diluent and serially diluted ¼ for n=8 points. After manual removal of blocking solution with a multichannel pipette, plasma titrations were added with one donor per column of the plate and incubated for 2 hours. The arrays were washed 3x 300 μL/well on the plate washer before adding 100 μL/well of a 1/5000 cocktail of biotinylated secondary polyclonal goat antibodies (Southern Biotech) directed to the human kappa and lambda light chains in assay diluent. After 1 hour of incubation and 3x 300 μL/well washing, a SA-HRP high sensitivity conjugate (Pierce) was added for 1 hour. After a final 3x 300 μL/well wash, residual buffer was manually removed with a pipette and 50 μL/well of sciCOLOR T2 added for 20 minutes.

### Conventional ELISA

The same high protein binding plates as used in the ELISA-Array were coated overnight at 4°C with 100 μL/well of the H1N1 A/Michigan/45/2015 strain of HA trimer in PBS, pH 7.2. The ELISA assay was performed in the same manner as the ELISA-Array with the exception of the development step. Plates were developed for 10 minutes with 50 μL/well soluble TMB substrate (KPL Sure Blue 1-component, SeraCare Life Sciences, Milford, MA) and stopped with 50 μL/well of TMB Stop solution (KPL).

## SUPPLEMENTARY MATERIAL

**Supplementary Figure S1. Optimization of printing parameters.** Initial tests of signal intensity using different Scienion formulation buffers (D01 vs. D11 vs. D12), protein stock concentrations (25 vs 100 vs 400 μg/mL), and number of drops (1 vs. 2 vs. 3). All spots are of goat anti-human Fc, made in triplicate, and detected using anti-kappa HRP. BLK indicates buffer background controls printed in the same array. Values in signal intensity are of human reference serum dilutions from dark to light green of 1/1,000, 1/5,000, 1/10,000, 1/100,000, and 1/1,000,000.

**Supplementary Table S1.** Intra-assay precision (n=3 plates) from all eight donor plasma samples, reference plasma, and 2 anti-Influenza mAbs on ELISA-array antigens.

**Supplementary Table S2.** Inter-assay precision (n=3 plates) from all eight donor plasma samples, reference plasma, and 2 anti-Influenza mAbs on ELISA-array antigens.

**Supplementary Table S3.** Inter-operator robustness (n=2 plates) from all eight donor plasma samples, reference plasma, and 2 anti-Influenza mAbs on ELISA-array antigens.

**Supplementary Table S4.** Stability of ELISA-Array plates at varying temperatures for 1 week (n=3 plates) from all eight donor plasma samples, reference plasma, and 2 anti-Influenza mAbs on ELISA-array antigens.

**Supplementary Table S5.** Stability of ELISA-Array plates under longer-term 4 °C storage (n=5 plates) from all eight donor plasma samples, reference plasma, and 2 anti-Influenza mAbs on ELISA-array antigens.

## AUTHOR CONTRIBUTIONS

KM and MS conceived and designed the study. EW, EC and KM carried out the experiments and performed data analysis. NF produced the recombinant viral antigens. MS coordinated and collected the donor plasma and provided the vaccine and virus materials. EW and KM wrote the manuscript. All the authors read and approved the final manuscript.

## ACKNOWLEDGEMENTS

The authors would like to thank Joshua Cantlon and Robert Kardish for their expert help with establishing the initial parameters and training for ELISA-array printing.

