## Supplementary figures and images for "Adaption of a Conventional ELISA to a 96-well ELISA-Array for Measuring the Antibody Responses to Influenza virus proteins, viruses and vaccines"

### Supplemental Figure 1

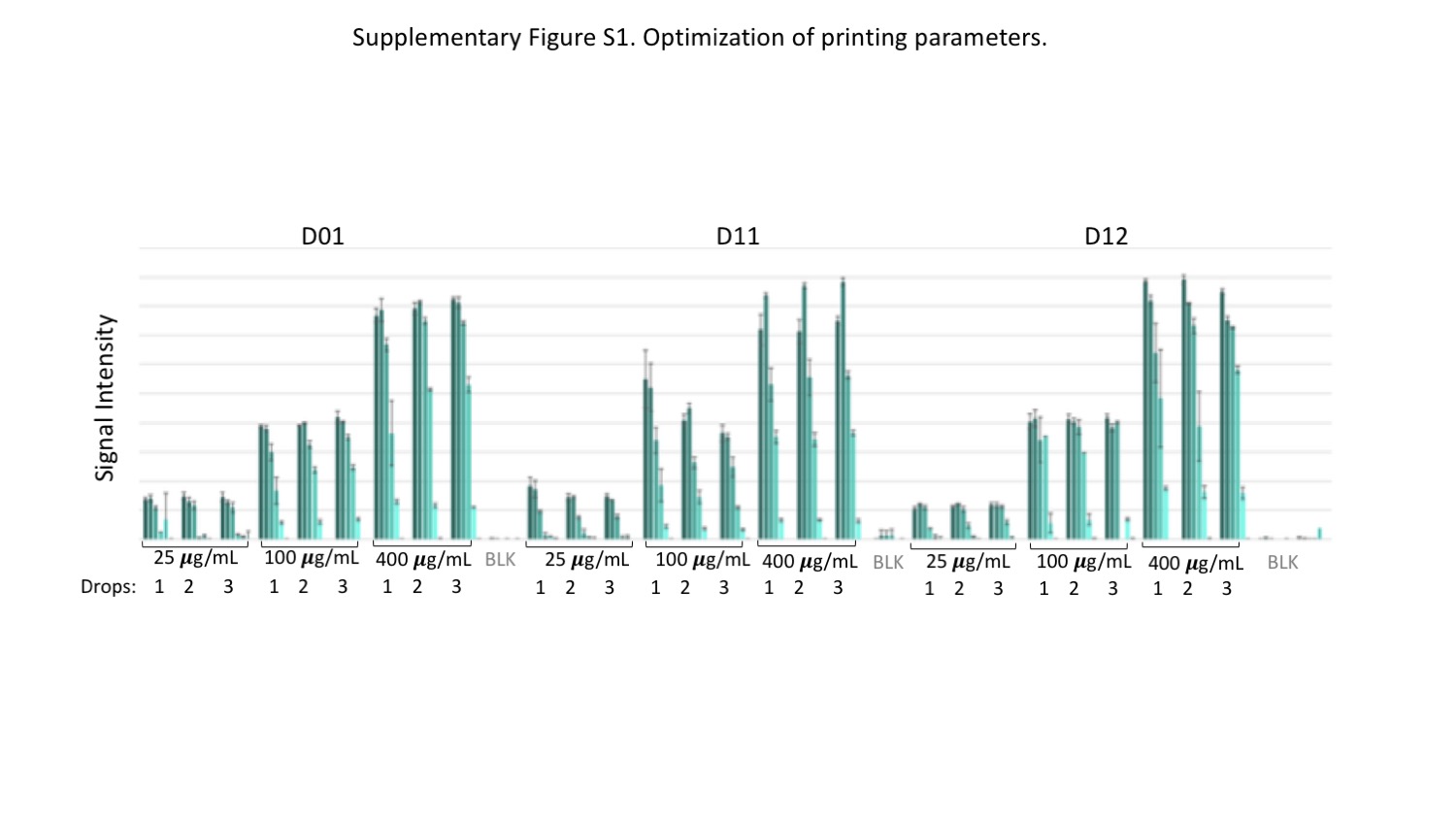
